# MYD88 mutations in clonal hematopoiesis promote inflammation and hematopoietic stem cell expansion

**DOI:** 10.1101/2025.06.19.660202

**Authors:** Jennifer Yeung, Sophia Y. Philbrook, Emma Uible, Lynn Lee, Kwangmin Choi, Puneet Agarwal, Courtnee A. Clough, Pravin Patel, Kathryn A. Wikenheiser-Brokamp, Kathleen Hueneman, Hans Christian Reinhardt, Tzyy-Jye Doong, Arletta Lozanski, Gerard Lozanski, Tamanna Haque, Erin Hertlein, John C. Byrd, Daniel T. Starczynowski

**Affiliations:** Division of Experimental Hematology and Cancer Biology, Cincinnati Children’s Hospital, Cincinnati, OH; Division of Oncology, Cincinnati Children’s Hospital, Cincinnati, OH; Department of Pathology and Laboratory Medicine, Hospital of the University of Pennsylvania, Philadelphia, PA; Division of Pathology and Laboratory Medicine, Cincinnati Children’s Hospital, Cincinnati, OH; Department of Hematology and Stem Cell Transplantation, West German Cancer Center, Partner Site Essen, University Hospital Essen, Essen, Germany; Division of Hematology, Department of Internal Medicine, The Ohio State University, Columbus, OH; Department of Pathology, The Ohio State University, Columbus, OH; Division of Leukemia, Memorial Sloan Kettering Cancer Center, New York, NY; Department of Internal Medicine, University of Cincinnati, Cincinnati, OH; University of Cincinnati Cancer Center, Cincinnati, USA; Department of Cancer Biology, University of Cincinnati, Cincinnati, OH; Department of Pediatrics, University of Cincinnati, Cincinnati, OH

## Abstract

Clonal hematopoiesis of indeterminate potential (CHIP) is characterized by expansion of mutant hematopoietic stem and progenitor cells (HSPCs) and an increased risk of chronic diseases and cancers. While mutations in *DNMT3A*, *TET2*, and *ASXL1* are common in CHIP, the contribution of less frequent gene mutations is not well understood. Here, we report *MYD88* mutations, including lymphoma-associated and novel variants in blood cells of the general population and newly diagnosed solid cancer patients. *MYD88* CHIP mutations in HSPCs activate NF-κB, indicating a gain-of-function activity. Modeling *MYD88* CHIP in mice, *Myd88*^L252P^ (equivalent of human L265P) expression resulted in a competitive fitness advantage of HSPCs. *Myd88*^L252P^ HSPCs exhibit a myeloid cell bias and inflammation, leading to hematologic disease. Single-cell RNA sequencing indicated that *Myd88*^L252P^ expands distinct hematopoietic and immune cell clusters and activates immune-related pathways in HSPCs. An IRAK1/4 inhibitor suppressed MYD88-dependent NF-κB activation and reversed *Myd88*^L252P^ cell expansion. Overall, *MYD88* mutations contribute to CHIP by inducing innate immune pathways in HSPCs and inflammatory disease.

## Introduction

Clonal hematopoiesis is an age-associated process in which the hematopoietic system is disproportionately derived from a single hematopoietic stem cell (HSC) clone^1^. This phenomenon, known as clonal hematopoiesis of indeterminate potential (CHIP), is defined by somatic mutations in known blood cancer-associated genes with a variant allele fraction (VAF) of 2% or greater in individuals without evidence of hematologic malignancy^1,2^. Individuals with CHIP are at an increased risk of developing a variety of age-related chronic diseases, including hematologic malignancies, such as myelodysplastic syndromes (MDS) or acute myeloid leukemia (AML), as well as cardiovascular diseases^1,2^. Depending on the cancer-associated mutation, certain individuals could also be at risk for lymphoid malignancies. CHIP-associated mutations confer a selective advantage, leading to an expanded clone. In the context of CHIP, mutations in a few genes, such as *DNMT3A, TET2*, and *ASXL1*, account for the majority of cases^3–6^. Less frequently mutated, but still prevalent genes, such as *TP53* and *U2AF1*, are associated with a higher risk of malignant transformation and adverse outcome^7,8^. This suggests that certain CHIP mutations confer varying risks of disease progression. Moreover, other genes, though less frequently mutated, collectively contribute to a significant proportion of CHIP cases^9^. The contribution and pathogenesis of these less common, but highly prevalent, gene mutations to CHIP remain poorly understood.

Dysregulated innate immune and inflammatory signaling, and alterations in the immune microenvironment, are critical mechanisms underlying clonal hematopoiesis^10^. Low-grade, chronic inflammation and other microenvironmental factors favor the expansion of HSCs with CHIP mutations over normal HSCs^10^. HSPCs with CHIP-related mutations also exhibit differential activation of innate immune and inflammatory pathways compared to age-matched HSPCs^10–13^. This dysregulation in immune-related pathways further contributes to the selective advantage of mutant HSCs and immune effector cells in CHIP. The Toll-like receptor (TLR) and Interleukin 1 receptor (IL1R) superfamily plays a key role in CHIP^4,10^. Upon activation, the Myeloid Differentiation Primary Response 88 (MYD88) adaptor, recruits IL-1 receptor-associated kinases (IRAKs) and TRAF6, leading to the activation of NF-κB and other transcription factors. Dysregulation of IRAK1, IRAK4, and TRAF6 has been implicated in the competitive fitness advantage of CHIP mutant HSPCs and altered immune cell activation^14–16^. *TET2* mutations in CHIP are associated with clonal expansion of HSPCs and increased inflammatory signaling via IRAK-TRAF6, which can drive the development of atherosclerosis^6^. Similar effects have been observed with TNFα-induced activation of NF-κB and the expansion of *DNMT3a*-mutant HSPCs.^17,18^ Moreover, patients with solid tumors who harbor tumor-infiltrating CHIP mutant cells have a higher risk of tumor recurrence and mortality compared to patients without CHIP, likely due to increased local inflammation^19^. To mitigate this inflammatory environment, inhibition of IL-1β has been shown to suppress mutant CHIP cells, partially restore normal hematopoiesis, and reduce cardiovascular disease in mouse models and early-stage clinical trials^20,21^.

Mutations in *DNMT3a, TET2*, and *ASXL1* constitute approximately 70% of CHIP^1,22–25^. It remains unclear whether the less common CHIP mutations similarly contribute to increased inflammation-related comorbidities, cardiovascular disease, or an elevated risk of hematologic malignancies. Understanding the role of less common CHIP mutations and their impact on the competitive advantage of mutant HSCPs and altered immune cells offers valuable insights into potential therapeutic targets in affected individuals. Our interest was piqued by recurrent mutations in *MYD88*, which have been identified in multiple clonal hematopoiesis studies, but often attributed to lymphoid origin monoclonal B-cell lymphocytosis. Mutations in *MYD88*, such as the amino acid substitution of Leucine 265 to Proline (L265P), are commonly found in lymphoid malignancies, such as Waldenstrom’s macroglobulinemia and chronic lymphocytic leukemia^26^. These gain-of-function mutations result in the constitutive activation of NF-κB, promoting cell survival and proliferation independent of external stimuli. MYD88 mutations have also been identified within CD34+ HSPCs at a preneoplastic stage in human lymphomagenesis^27^. Collectively, these findings suggest that *MYD88* mutations occur in multi-lineage HSPCs, confer a competitive fitness advantage associated with CHIP, and increase the risk of certain malignancies.

## Results

### Recurrent CHIP-associated *MYD88* mutations

To determine the distribution of *MYD88* mutations in healthy adult individuals, we analyzed published sequencing data of 200,618 individuals aged 40-70 years from the UK Biobank (UKBB)^28^. The genes with the highest prevalence of candidate somatic nonsense mutations were *DNMT3A, TET2*, and *ASXL1*. Genes, such as *ZBTB33*, *YLPM1, ZNF318, JAK2*, and *SRCAP*, with lower frequency mutations in CHIP have been shown to also confer a competitive advantage to HSPCs^29–31^. The number of *MYD88* mutations in this cohort is in a similar range as the lower frequency CHIP mutations (**Figure 1A**). To examine the frequency of *MYD88* mutations in additional published CHIP cohorts, we analyzed 6 additional independent studies, consisting mostly of individuals of European ancestry (**Supplemental Tables 1-2**). The studies included both healthy individuals and those diagnosed with cytopenias or cancer. Collectively, 1-27 *MYD88* CHIP mutations were reported, representing ∼0.5-2% of identified CHIP individuals (**Figure 1B**). To confirm these observations in an independent cohort, we performed targeted error-corrected sequencing of mononuclear cells from newly diagnosed solid tumor cancer patients (**Supplemental Table 3**). In this cohort, we observed a higher than expected proportion (∼4%) of individuals harboring *MYD88* CHIP mutations: 10 mutations in 9 individuals out of 218 confirmed CHIP cases among 454 cancer patients sequenced (**Figure 1C**). The validation cohort consists entirely of individuals from the rural Appalachian region, all of whom are Caucasian, with newly diagnosed solid tumors, which are geographically underrepresented in prior genomic studies. The variant allele frequency of *MYD88* ranged from 2-36% (**Supplemental Table 3**). By comparison, the TCGA and MSK cohorts have a *MYD88* mutation frequency of 0.32% and 0.14%, respectively (**Supplemental Table 2**). The mutations in *MYD88* were missense variants clustered in the death domain and TIR domain (**Figure 1C**). Among all *MYD88* CHIP variants, we noted a high frequency of the oncogenic mutation at L265P within the TIR domain commonly found in B-cell lymphomas. In the Appalachian cohort, we also identified mutations at A45G, T71I, S85P within the death domain, R140Q in the linker region, and C203R within the TIR domain (**Figure 1C**). Although the C203R variant is located within a GC-rich region, which results in lower quality reads, an examination of the Genome Aggregation Database revealed that MYD88 CHIP variants have been reported at or near this residue within the TIR domain in other cohorts. This suggests that mutations within the TIR domain, beyond L265P, can influence MYD88-dependent signaling^32^ (**Supplemental Table 4**). These mutations were not previously reported among MYD88 variants detected in human lymphoma^33^. Moreover, mutations in *MYD88* are not reported in MDS and AML^34^. Future studies should examine a larger and more genetically diverse cohort to better understand the broader relevance of these findings. Furthermore, since we detected MYD88 mutations in blood cells from tumor patients, the potential contribution of MYD88-mutant CHIP to solid tumor progression warrants further investigation. Overall, the *MYD88* mutations identified in CHIP, which appear to be underreported, were not associated with a B cell malignancy diagnosis, suggesting they may have a distinct role in CHIP.

**Figure 1.**
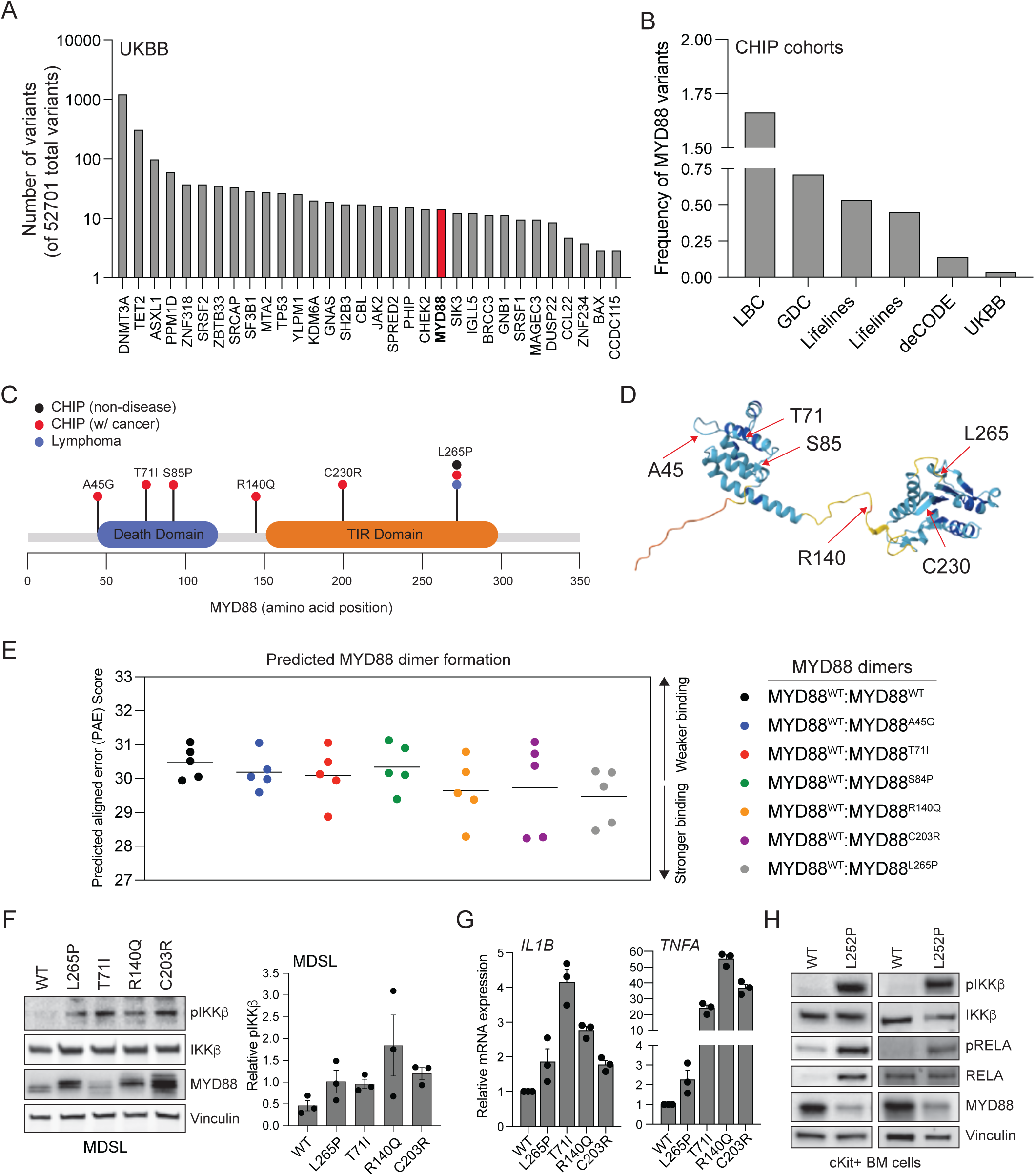
CHIP-associated MYD88 mutations confer a gain-of-function. **(A)** The distribution of identified CHIP-associated somatic mutations ranked by frequency in UK Biobank (Bernstein, Nature Genetics, 2024). (**B**) The frequency of MYD88 mutations identified in the indicated population cohorts. (C) Lollipop diagram of CHIP-associated mutations identified in MYD88. CHIP-associated mutations were identified in individuals with no known disease phenotype (black), a solid tumor cancer diagnosis ^15^, or lymphoma (blue). (**D**) Prediction of human MYD88 protein structure using AlphaFold^72^. Highlighted residues are implicated in CHIP: A45, T71, S85, R140, C203, and L265. **(E)** Prediction of protein-protein interactions using AlphaFold 3. Predicted aligned error ^29^ score of the indicated human MYD88 dimers: WT and WT, WT and L265P, WT and T71I, WT and R140Q, WT and A45G, WT and S85P, and WT and C203R. (**F**) Representative immunoblots of MDSL MYD88 knockout (KO) cells expressing WT and CHIP-related MYD88 mutants from 3 independent experiments. Quantification of phosphorylated IKKβ (relative to total IKKβ) immunoblots (right panel). **(G)** Quantitative RT-PCR (RT-PCR) analysis of NF-κB target genes (*TNFA* and *IL1B*) in THP1 MYD88 KO cells transduced with lentiviral vectors encoding WT MYD88 or the CHIP mutations (L265P, T71I, R140Q, and C203R) Data from 3 independent biological replicates. (**H**) Immunoblots of cKit+ enriched BM cells from Myd88^WT^ and Myd88^L252P^ mice treated with 1 μM 4-hydroxytamoxifen (4-OHT) for 72 hours to induce Cre recombination.

### *MYD88* mutations result in NF-κB activation

Previous studies revealed that the L265P mutation results in spontaneous MYD88 assembly and constitutive NF-κB signaling in B cells without receptor activation^35^. Structural modeling using AlphaFold3 revealed that the CHIP mutations are clustered in key oligomerization regions of MYD88 (**Figure 1D**). Next, we aimed to gain insight into whether CHIP mutations in MYD88 affect oligomer formation. To assess this, we utilized predicted aligned error^29^, a measure of AlphaFold3’s confidence in the relative positioning of two residues within the predicted protein structure^36^. Comparisons between WT-WT MYD88 protein dimers and WT-mutant dimers revealed a lower mean PAE score for the WT-mutant MYD88 dimers (**Figure 1E**, **Supplemental Figure 1A**). This analysis predicts that all MYD88 mutants have a structural orientation favoring a stable dimer formation. To determine whether MYD88 CHIP mutations result in constitutive NF-κB signaling in hematopoietic cells, we generated isogenic cells expressing the individual *MYD88* CHIP mutations. We selected CHIP mutants that were recurrent and with the highest variant allele frequency: T71I, R140Q, C203R, and L265P. Endogenous MYD88 was deleted in myeloid-derived THP1 and MDSL cells using CRISPR/Cas9, and then the MYD88^KO^ cells were transduced with vectors encoding either WT MYD88 or the CHIP mutants. RNA sequencing revealed that MYD88 CHIP mutants lead to increased, yet distinct and variable, expression of NF-κB target genes, as compared to WT MYD88-expressing cells (**Supplemental Figure 1B**). These differences in NF-κB activation are expected, as individual CHIP-associated mutations may variably affect MYD88 signaling complex assembly, downstream effector recruitment, or stability. Consistent with findings in B-cell malignancies^35^, expression of MYD88 L265P in myeloid-derived cells resulted in constitutive activation of NF-κB, as indicated by increased phosphorylation of IKKβ, independent of TLR activation, as compared to WT MYD88 (**Figure 1F,G**). The other CHIP-associated MYD88 mutants - T71I, R140Q, C203R - also promoted activation of IKKβ, although to variable extents (**Figure 1F,G**). Increased expression of *TNFA* and *IL1B*, two canonical NF-κB target genes, further confirmed gain-of-function activity of MYD88 CHIP mutants (**Figure 1G**). Although the MYD88 CHIP mutants exhibit elevated baseline NF-κB activation, the addition of IL-1β did not result in enhanced signaling (**Supplemental Figure 1C**). Together, these results indicate that CHIP-associated MYD88 mutations confer gain-of-function activity, leading to constitutive, ligand-independent activation of the NF-κB pathway.

Although *MYD88* L265P results in constitutive NF-κB signaling in B cells^35,37^, it remains unclear whether this mutation similarly activates NF-κB in HSPCs. Since *MYD88* L265P is the most common MYD88 CHIP mutation, we investigated its role in HSPCs by characterizing a conditional Myd88 L252P allele (Myd88^L252P^) expressed from the endogenous locus following inducible Cre-mediated recombination^37^. The murine Myd88 L252P corresponds to the orthologous position of human MYD88 L265P. Myd88^L252P^ mice were crossed to *Rosa26^CreERT^*^2^ mice to allow for recombination following tamoxifen treatment^38^. We focused on characterizing the homozygous Myd88^L252P^ mice, as there is evidence of bi-allelic *MYD88* mutations in patients^27,39^, and the heterozygous expression of the mutant allele has been shown to elicit a modest effect on NF-κB activation in mouse cells^40^. To confirm Cre-inducible Myd88 L252P expression, we isolated BM cells from littermates WT (*Myd88^WT/WT^;RosaCreERT2*) and Myd88^L252P/L252P^ (*Myd88^L252P^;RosaCreERT2*) mice. 4-hydroxytamoxifen (4-OHT) treatment of Myd88^L252P^ BM cells resulted in Cre-induced recombination of the Myd88 locus, resulting in expression of *Myd88^L252P^*-mutant mRNA (**Supplemental Figure 2A**). 4-OHT-induced expression of *Myd88^L252P^* in cKit+ BM HSPCs resulted in IKKβ and RelA/p65 phosphorylation, an indication of NF-κB activation (**Figure 1H**). In contrast, WT HSPCs did not exhibit evidence of baseline NF-κB activation, suggesting that *MYD88* CHIP mutations are sufficient to induce NF-κB activation in HSPCs.

### *MYD88* mutations result in a competitive fitness advantage

Expression of CHIP mutations in murine HSPCs can result in increased serial colony replating, an indicator of enhanced self-renewal potential. To determine whether *MYD88* CHIP mutations have a cell-intrinsic effect on HSPCs, we assessed the colony-forming potential of HSPCs (cKit+) isolated from WT and *Myd88^L252P/L252P^* (herein referred to as *Myd88^L252P^*) mice in methylcellulose. HSPCs from *Myd88^L252P^* mice formed a greater number of colonies, predominantly resulting from increased myeloid-lineage forming colonies, as compared to WT HSPCs (**Figure 2A**). Moreover, *Myd88^L252P^* HSPCs exhibited an increase in colony replating potential for up to 5 serial platings as compared to WT HSPCs (**Figure 2B,C**). The increase in colony replating observed in *Myd88^L252P^* HSPCs is comparable to more common CHIP variants^41–48^.

**Figure 2.**
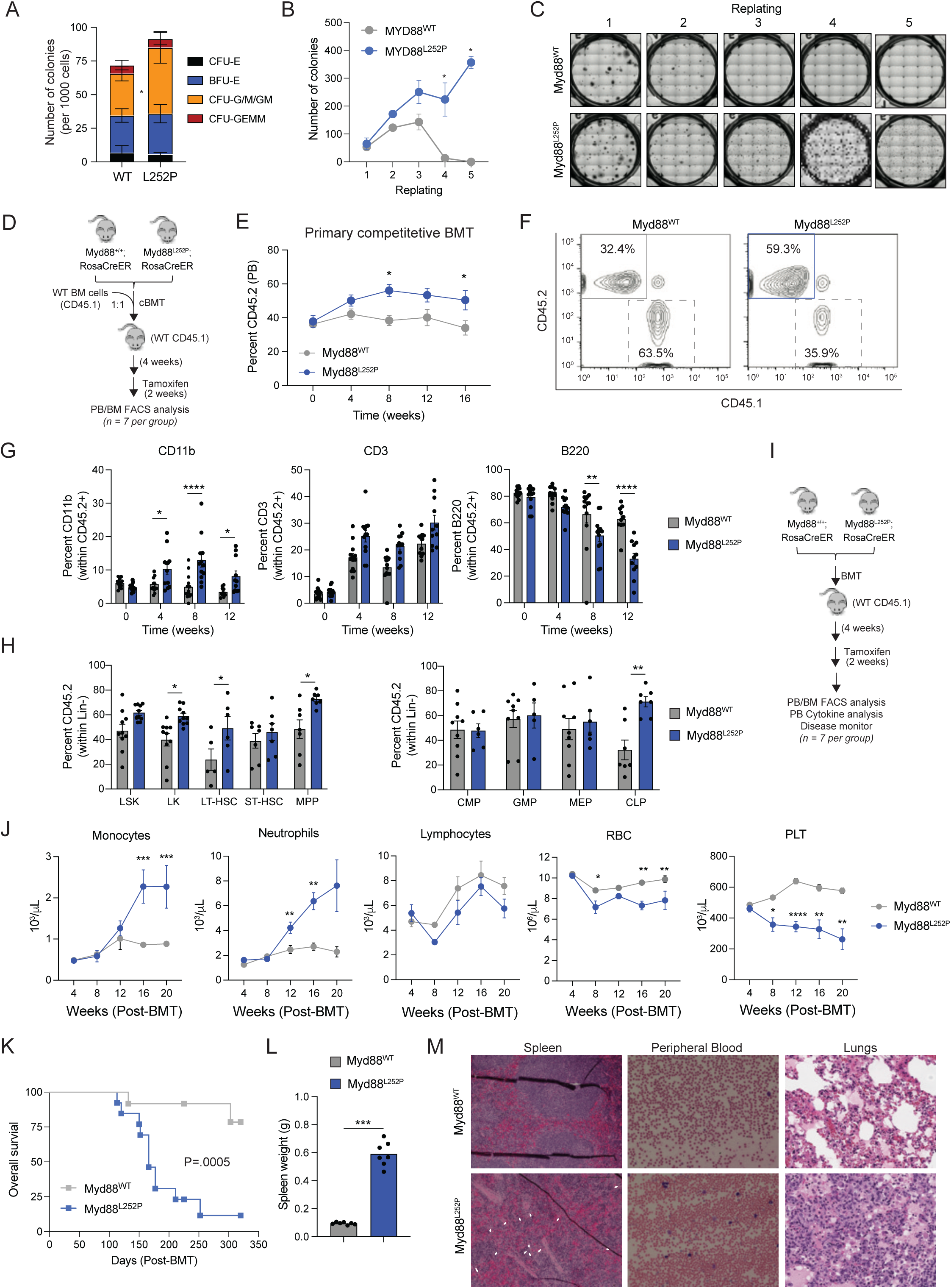
Myd88^L252P^ confers an HSPC competitive advantage, myeloid-biased hematopoiesis, and leads to an immune-related hematologic disease. (**A**) 4-OHT (1 μM) treated cKit+ enriched BM cells (n = 1000) from Myd88^WT^ and Myd88^L252P^ mice were plated in methylcellulose and assessed for colony formation (n = 3 per group from biological replicates): erythroid progenitor cells (BFU-E), granulocyte-macrophage progenitors (CFU-G/M/GM), and multi-potential granulocyte, erythroid, macrophage, megakaryocyte progenitor cells (CFU-GEMM). (**B**) Serial colony replating potential of cKit-enriched BM cells from MYD88^WT^ and MYD88^L252P^ mice in methylcellulose. Error bars represent the SEM (n = 3 per group from biological replicates). (**C**) Representative images of colonies from panel B. (C) Experimental overview of competitive BM transplants (cBMT). (**E**) Summary of donor-derived Myd88^WT^ and Myd88^L252P^ PB proportions (CD45.2) from the cBMT recipient mice at the indicated time points (n = 14-15 mice per group). (**F**) Representative flow cytometry plots of donor-derived CD45.1 (WT) or CD45.2 (Myd88^WT^ or Myd88^L252P^) PB cells from primary cBMTs. (**G**) Proportions of donor-derived CD45.2 populations from the PB of primary cBMTs: myeloid (CD11b^+^), T (CD3^+^), and B (B220^+^) cells at the indicated time points (n= 11-13 mice per group). Error bars represent SEM (n = 11-13 per group). (**H**) Proportions of donor-derived CD45.2 populations from the BM of primary cBMTs at 16 weeks (n = 7-10 mice per group): LK (Lin^-^cKit^+^Sca1^-^), LSK (Lin^-^ckit^+^Sca1^+^), LT-HSC (LSK CD150^+^CD48^-^), ST-HSC (LSK CD150^-^CD48^-^), MPP (LSK CD150^-^CD48^+^), CMP (LK CD34^+^16/32^-^), MEP (LK CD34^-^CD16/32^-^), GMP (LK CD34^+^CD16/32^+^), CLP (Lin^-^ckit^-^Sca1^lo^CD127^+^CD135^+^). (**I**) Experimental overview of non-competitive BM transplants (BMT). (**J**) Complete PB counts of MYD88^WT^ and MYD88^L252P^ at the indicated time points post BM transplantation and tamoxifen administration. Error bars represent SEM (n = 7-8 mice per group). (**K**) Kaplan-Meier survival curves for recipient mice transplanted with MYD88^WT^ (n = 12) and MYD88^L252P^ BM cells (n = 13 mice per group). Data from 2 independent biological replicates. (**L**) Weight of spleen isolated from recipient mice transplanted with Myd88^WT^ and Myd88^L252P^ BM cells. Error bars represent SEM (n = 7 mice per group). (**M**) Representative images of spleen sections (Hematoxylin and Eosin), PB smears (Wright-Giemsa), and lungs (Hematoxylin and Eosin) from recipient mice transplanted with Myd88^WT^ and Myd88^L252P^ BM cells. White arrows denote infiltrates of monocytes or neutrophils in the spleen. Scale bars of H&E and Wright-Giemsa images represent 50 μm. Error bars represent SEM. Significance was determined with a Student’s t-test for two groups or ANOVA for multiple groups (*, P < 0.05; **, P < 0.01; ***, P < 0.001).

To evaluate the impact of Myd88 mutations on clonal hematopoiesis, we performed *in vivo* competitive repopulation assays using BM cells from *Myd88^WT^;Rosa26CreERT2* (hereafter WT) and *Myd88^L252P^;Rosa26CreERT2* (hereafter *Myd88^L252P^*) mice transplanted with equal numbers of WT competitor cells into lethally irradiated recipient mice. Following engraftment, mice were administered with tamoxifen to induce expression of the *Myd88* mutant allele and then analyzed every 4 weeks (**Figure 2D**). Donor-derived *Myd88^L252P^* cells outcompeted WT cells over time in the peripheral blood (**Figure 2E,F**). In the competitive BM transplantation, *Myd88^L252P^* expression contributed to an expansion of donor-derived myeloid cells (CD11b) and a relative suppression of B cells (B220) (**Figure 2G**). This contrasts with the B cell-restricted CD19-Cre *Myd88^L252P^* model, which shows no changes in the B220⁺ population in the BM^40^. We observed an expansion of *Myd88^L252P^* immature neutrophils (CD11b^+^Gr1^Low^)(**Supplementary Figure 2B,C**). Since *Myd88^L252P^* expression resulted in the expansion of myeloid cells in the PB, we next examined the HSPC proportions in the BM of these mice. The proportions of donor-derived *Myd88^L252P^* long-term HSCs (49.05 vs 23.73%), multi-potent progenitors (72.74 vs 44.5%), and common lymphoid progenitors (CLP)(71.74 vs 32.26%) were significantly increased as compared to WT donor-derived populations (**Figure 2H** and **Supplemental Figure 2D**). We postulate that the increase in donor-derived *Myd88^L252P^* CLPs is a compensatory response to the reduced proportions of mature B cells and may eventually contribute to the increased expansion of B cells as observed in B cell malignancies. These observations suggest that *MYD88* CHIP mutations promote myeloid-biased hematopoiesis, with a particular effect on monocytes and immature neutrophils, and a competitive fitness advantage of HSPCs. Moreover, the ability of MYD88 L252P to promote both lymphoid and myeloid cell alterations highlights the versatility of *MYD88* mutations and their capacity to drive clonal expansion across different hematopoietic lineages even in the absence of eventual progression to myeloid malignancy.

### *MYD88* mutations result in an inflammatory disease

*MYD88* CHIP mutations are associated with a heightened risk of lymphoid malignancies and increased all-cause mortality^49–51^. Moreover, *MYD88* CHIP mutations were observed in certain individuals with a VAF exceeding >30%^52^, suggesting that the majority of the BM and blood cells contain the mutation. To determine whether *MYD88* CHIP mutations can lead to increased mortality and lymphoid malignancy development, we next evaluated the effect of *Myd88^L252P^* expression on hematopoiesis in a non-competitive BM transplantation model (**Figure 2I**). Mice transplanted with *Myd88^L252P^* BM cells exhibited increased neutrophils (P < 0.01) and monocytes (P < 0.05), resembling a myeloproliferative disorder, and reduced red blood cells (P < 0.01) and platelets (P < 0.001) 12 weeks post-transplantation as compared to mice transplanted with WT BM cells (**Figure 2J**). No significant changes in lymphocytes were observed in mice transplanted with *Myd88^L252P^* BM cells. Interestingly, CH individuals with unexplained anemia have been found to harbor MYD88 mutations^53^, supporting the observation of reduced red blood cells in the *Myd88^L252P^* mice. Recipient mice transplanted with *Myd88^L252P^* BM cells had a shorter overall survival as compared to control mice (**Figure 2K**). Nearly all (∼90%) *Myd88^L252P^* recipient mice succumbed to disease within 1 year following BM transplantation. Moribund *Myd88^L252P^* recipient mice exhibited enlarged spleens and disruption of spleen architecture, including extramedullary hematopoiesis (**Figure 2L,M, Supplemental Figure 2E**). The spleens of *Myd88^L252P^* recipient mice had infiltrates of monocytes and neutrophils, features not observed in B cell-specific expression of Myd88^L252P^. The liver of *Myd88^L252P^* recipient mice also displayed mild periportal mononuclear cell infiltrates with hepatocyte injury and occasional apoptotic hepatocytes (**Supplemental Figure 2E**). Additionally, the lungs of *Myd88^L252P^* recipient mice harbored peribronchiolar and perivascular inflammation, characterized by lymphocyte infiltration, which likely contributes to the observed morbidity and reduced overall survival in these mutant mice (**Figure 2M**). B cell expression of *Myd88^L252P^* resulted in the expansion of splenic small mature lymphocytes or large blastoid cells in the liver^37^. However, examination of the BM and PB in *Myd88^L252P^* recipient mice did not reveal a lymphoid or myeloid malignancy, suggesting that *MYD88* CHIP mutations do not result in overt hematologic cancer. Conversely, B cell-specific *MYD88^L252P^* expression harbored lymphoma cells or activated lymphocytes^37^. These findings revealed that expression of *Myd88^L252P^* in hematopoietic cells results in distinct myeloid and lymphoid phenotypes.

Since we observed significant changes in mature immune cells and a non-malignant hematologic disease in *Myd88^L252P^* recipient mice, we posited that the *MYD88* CHIP mutations contribute to disease due to chronic NF-κB-induced cytokine production. Thus, to determine whether *Myd88^L252P^* recipient mice exhibit altered cytokine expression, we collected plasma from the BM of the non-competitive BM transplantation models and profiled 32 murine cytokines (**Figure 2I**). Peripheral blood plasma from *Myd88^L252P^* recipient mice had a broad increase in multiple cytokines, such as IL-6, TNFα, G-CSF, MIG, MIP1α, and IFNγ (**Figure 3A,B, Supplemental Table 5**). Chronic inflammation can enhance myeloid expansion and selectively expand the mutant clones at the expense of suppressing normal HSCs, indicating that persistent inflammatory signaling is a critical driver of CHIP progression^11,54,55^. Collectively, these findings suggest that *MYD88* CHIP promotes an inflammatory milieu, which can lead to increased mortality.

**Figure 3.**
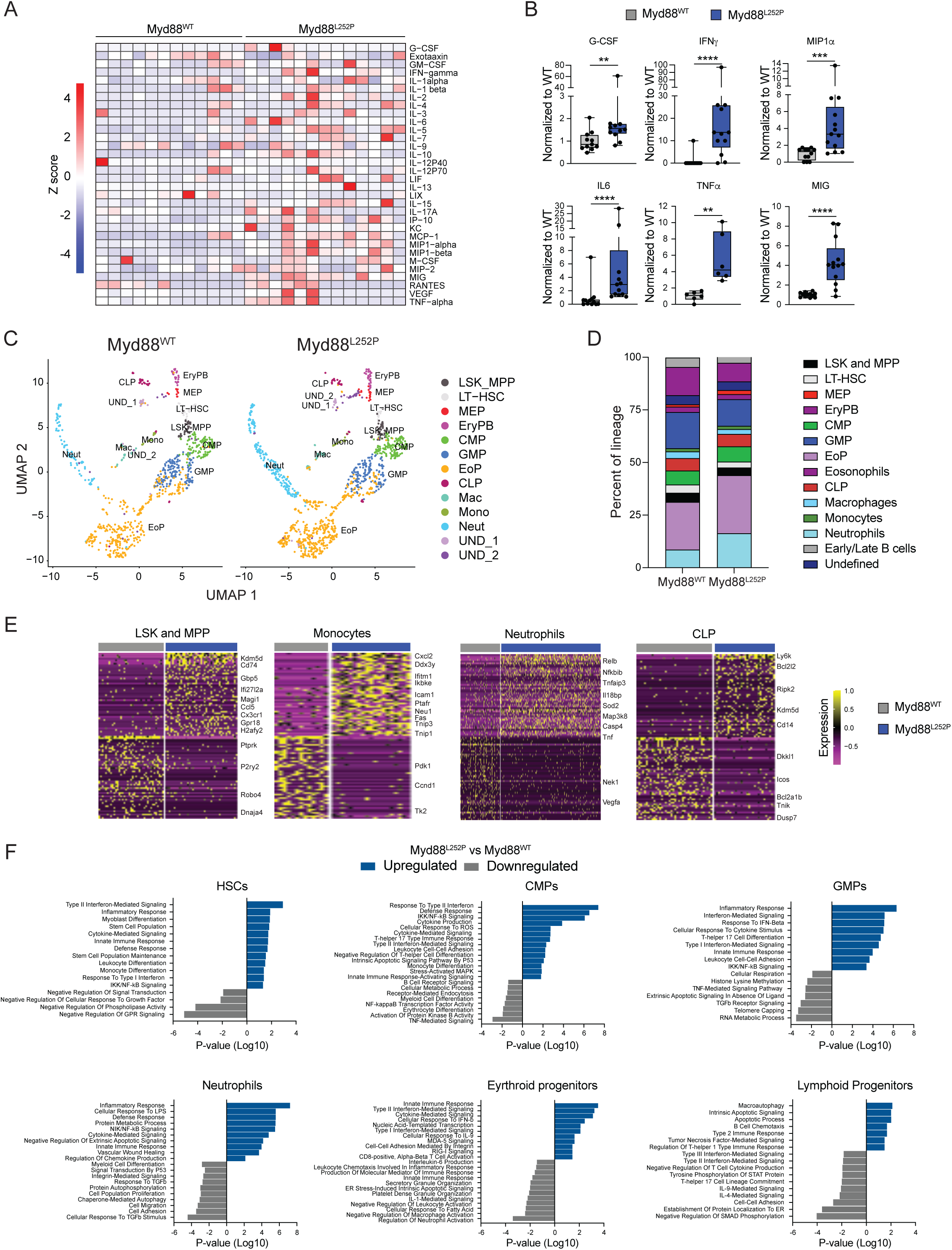
scRNA-seq of Myd88^L252P^ BM HSPCs reveals broad immune and inflammatory signaling dysregulation. (**A**) PB plasma cytokines were evaluated after 20 weeks in recipient mice transplanted with Myd88^WT^ (n = 12) and Myd88^L252P^ (n = 13) BM cells. A heatmap of plasma cytokine levels is shown and color-coded according to Z score (high, red; middle, orange; low, blue). (**B**) Representative cytokines from Myd88^L252P^ PB are normalized to Myd88^WT^ PB. Significance was determined with a Student’s t-test for two groups or ANOVA for multiple groups (*, P < 0.05; **, P < 0.01; ***, P < 0.001). (**C**) BM cells were harvested from recipient mice transplanted with Myd88^WT^ and Myd88^L252P^ BM cells (n = 2 mice per group) after 12 weeks, and cKit+ cells were enriched by magnetic selection. Barcoded cells were pooled and analyzed by 10X single-cell RNA (scRNA) sequencing. UMAP of 900 Myd88^WT^ and 1195 Myd88^L252P^ cells representing 13 clusters. **(D)** Percentage of each cell cluster from the scRNA-seq analysis of 900 Myd88^WT^ or 1195 Myd88^L252P^ cells. **(E)** Heatmap of the top 50 differentially expressed genes in individual Myd88^WT^ or Myd88^L252P^ cells within the indicated clusters. **(F)** Gene ontogeny pathway analysis of the indicated clusters. All error bars represent SEM. Significance was determined with a Student’s t-test for two groups or ANOVA for multiple groups (*, P < 0.05; **, P < 0.01; ***, P < 0.001).

### scRNA-seq reveals insights into *MYD88*-mutant CHIP

To understand how *Myd88^L252P^* HSPCs result in a competitive fitness advantage, myeloid expansion, and inflammatory pathology, we performed single-cell RNA sequencing (scRNA-seq) on 10,000 sorted cKit+ BM cells isolated from recipient mice engraftment with either WT or *Myd88^L252P^* BM cells 16 weeks following engraftment (as in Figure 2I). We identified 13 major cell lineage clusters in both WT and Myd88^L252P^ cKit+ BM cells (**Figure 3C**). Consistent with our flow cytometry analysis, *Myd88^L252P^* cKit+ BM cells showed an expansion of neutrophil, common myeloid progenitor (CMP), and megakaryocyte-erythroid progenitor (MEP) clusters compared to the WT BM cells (**Figure 3D**). Moreover, clusters consisting of granulocyte monocyte progenitors (GMP), eosinophils, and B cells were reduced in the *Myd88^L252P^* BM as compared to WT mice (**Figure 3D**). We then sought to delineate the developmental trajectory between the immature HSPCs and the committed progenitors by a pseudotime analysis^56^. The trajectory of *Myd88^L252P^* cells towards neutrophils was more prominent as compared to WT cells (**Supplemental Figure 3A**). These findings indicate that the MYD88 mutation promotes the myeloid commitment of BM HSPCs and expands immature neutrophils.

To identify mechanisms by which *Myd88^L252P^* promotes long-term fitness advantage, we focused on the HSPC clusters. Our findings revealed a substantial increase in upregulated differentially expressed genes (DEGs) in *Myd88^L252P^* clusters containing HSPCs (LSK/MPP), monocytes, neutrophils, and CLP compared to WT clusters (**Figure 3E** and **Supplemental Table 6**). Gene Ontology (GO) analyses highlighted enrichment for Type I interferon, innate immune response, myeloid differentiation, and stem cell maintenance programs in *Myd88^L252P^* HSPCs as compared to WT HSPCs (**Figure 3F**). Similarly, *Myd88^L252P^* clusters containing monocytes and neutrophils had a significant increase in the number of DEGs as compared to WT cells (**Figure 3E**). Within the myeloid cell clusters, *Myd88^L252P^* cells demonstrated increased inflammatory genes (**Figure 3F**). *Myd88^L252P^* clusters containing lymphocyte progenitors exhibited enrichment of autophagy and cell death programs (**Figure 3F**). While common CHIP mutations typically drive myeloid-biased hematopoiesis, our findings reveal that *MYD88* mutations may have a more complex role, influencing both myeloid and lymphoid programs.

### IRAK1/4 inhibition suppresses *MYD88-*mutant cells

The size of the mutant hematopoietic cell pool can predict the progression to hematological cancers and immune-related conditions, therefore, strategies to prevent the expansion of mutant clones are clinically desirable. Individuals with *MYD88* CHIP mutations are at an increased risk of hematological cancers and cardiopulmonary diseases^49–51^. As such, preventing the expansion of *MYD88* CHIP mutant clones, or other CHIP mutant clones exhibiting IRAK1/4 signaling, could be a feasible clinical intervention. IRAK4 inhibitors have been explored as potential therapies for lymphoid malignancies driven by *MYD88* mutations^16,57^. In preclinical studies, IRAK1/4 inhibition suppressed pre-leukemic and MDS/AML cells, suggesting that dysregulation of IRAK1/4 is a key driver of mutant HSPCs and a potential therapeutic target^14,16,58–61^. To determine whether MYD88-dependent signaling is responsible for the competitive advantage of *MYD88* CHIP mutant cells, we used an inhibitor (NCGC-1481, herein “IRAK1/4-inh”) to suppress IRAK1 and IRAK4, the proximal effectors of MYD88^62–64^. The IRAK1/4-inh effectively repressed NF-κB activation and innate immune (*Ifng*) and inflammatory (*Mip1a* and *Tnfa*) gene expression in *Myd88^L252P^* cKit+ BM cells, but not in *Myd88^WT^* cells (**Figure 4A, Supplemental Figure 3B**). Since *Myd88^L252P^* HSPCs acquire an enhanced colony-replating phenotype, we aimed to determine if this was due to increased innate immune signaling. Treatment with the IRAK1/4-inh did not affect the primary colony-forming potential of either *Myd88^WT^* or *Myd88^L252P^* cKit+ BM cells. However, it significantly reduced the secondary, tertiary, and quaternary colony-forming potential of *Myd88^L252P^* cKit+ cells, while having no effect on *Myd88^WT^* cells (**Figure 4B**).

**Figure 4.**
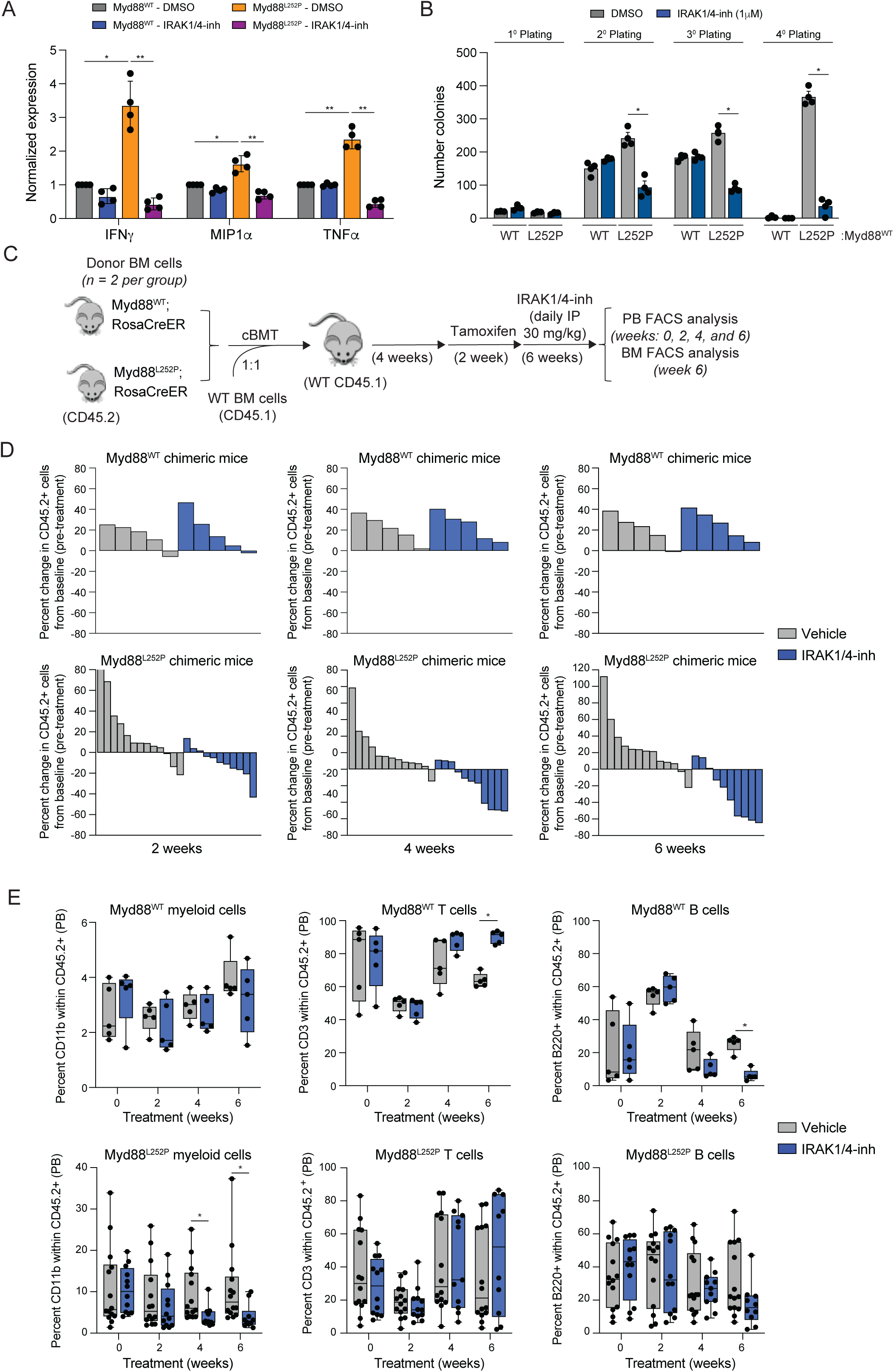
IRAK1/4 inhibition suppresses innate immune signaling, clonal expansion, and the competitive fitness advantage of Myd88^L252P^ HSPCs. (A) Quantitative RT-PCR (RT-PCR) analysis of the indicated cytokines genes in Myd88^WT^ and Myd88^L252P^ cKit+ BM cells incubated with 4-OHT to induce recombination for 72 hours followed by treatment with vehicle (DMSO) or 1 μM IRAK1/4-inhibitor (IRAK1/4-inh; NCGC-1481) for 24 hours (n = 3 per group from biological replicates). (B) cKit+ enriched BM cells incubated with 4-OHT (1 μM) were serially plated in methylcellulose and assessed for colony formation in the presence of vehicle (DMSO) or 1 μM IRAK1/4-inh (NCGC-1481) (n = 3 per group from biological replicates). (C) Experimental overview of competitive BM transplants (cBMT) following administration of the IRAK1/4-inh (NCGC-1481). Chimeric mice were administered daily with either vehicle control (PBS) or IRAK1/4-inh (30 mg/kg) via intraperitoneal (IP) injection for 6 weeks. FACS analysis of PB was performed pre- and post-administration at weeks 0, 2, 4, and 6. (D) Percent change (related to pre-treatment) in PB chimerism of individual donor-derived CD45.2 cells following administration of vehicle or IRAK1/4-inh at the indicated time points (n = 10 for Myd88^WT^; n = 26 for Myd88^L252P^). Data from 2 independent biological replicates. (E) Proportions of PB donor-derived myeloid (CD11b), T (CD3), and B (B220) cells within the CD45.2 populations of Myd88^WT^ (n = 10) and Myd88^L252P^ (n = 26) chimeric mice following treatment with vehicle or IRAK1/4-inh at the indicated time points. Data from 2 independent biological replicates. Error bars represent SEM. Significance was determined with a Student’s t-test for two groups or ANOVA for multiple groups (*, P < 0.05; **, P < 0.01; ***, P < 0.001).

We next performed an *in vivo* competitive repopulation assay using CD45.2 BM cells from *Myd88^WT^* or *Myd88^L252P^* mice, as above, by transplanting equal numbers of CD45.1 WT competitor cells into lethally irradiated recipient mice. Following engraftment (4 weeks post-transplant), mice were administered tamoxifen to induce expression of the *Myd88-* mutant allele. One week post tamoxifen administration, chimeric recipient mice were randomized and treated daily for 6 weeks with the IRAK1/4-inh or vehicle (PBS) and analyzed every 2 weeks for hematopoietic reconstitution (**Figure 4C**). As expected, donor-derived *Myd88^L252P^* cells in the vehicle group outcompeted WT cells (**Figure 4D**). Treatment of the *Myd88^L252P^* recipient mice with the IRAK1/4-inh suppressed donor-derived *Myd88^L252P^*, but not *Myd88^WT^*cells in the PB (**Figure 4D**). The IRAK1/4-inh also suppressed donor-derived *Myd88^L252P^* HSPCs in the BM, including MPPs, GMPs, and MEPs (**Supplemental Figure 3C**). Examination of the PB and BM following IRAK1/4 inhibitor treatment revealed that the reduced chimerism of donor-derived *Myd88^L252P^* cells was primarily due to a decrease in the proportions of CD11b myeloid cells in the PB (**Figure 4E**). IRAK1/4-inh had negligible effects on the chimerism of *Myd88^WT^* cells in the PB and HSPCs (**Figure 4D** and **Supplemental Figure 3C**). These findings suggest that the competitive advantage of MYD88-mutant CHIP cells is driven, at least in part, by IRAK1/4 signaling and can be reversed by small molecule inhibition. This highlights aberrant innate immune signaling as a mechanism underlying MYD88-mutant CHIP HSC expansion and supports IRAK1/4 inhibition as a potential therapeutic strategy.

## Discussion

Our findings reveal that MYD88 mutations, though underreported in CHIP, confer a cell-intrinsic competitive advantage by promoting innate immune activation and chronic inflammation. This establishes a direct mechanistic link between MYD88 signaling and clonal expansion. We show that recurrent MYD88 CHIP mutations, including L265P and other TIR/death domain variants, promote NF-κB activation. In vivo, MYD88-mutant HSPCs exhibited enhanced self-renewal capacity, myeloid-biased differentiation, and competitive repopulation potential. MYD88-mutant cells also induced inflammatory disease, mimicking CHIP-associated comorbidities. These effects occurred in the absence of overt hematologic malignancy, suggesting that MYD88 mutations can drive disease through inflammation alone. Gene expression profiling revealed a shift toward myeloid lineages and enrichment of inflammatory gene programs in MYD88-mutant HSPCs. These data highlight the broad transcriptional and lineage consequences of MYD88 signaling in CHIP. Finally, we show that IRAK1/4 inhibition selectively suppresses MYD88-mutant HSPCs, reducing their fitness and expansion without affecting WT cells. These results position the MYD88–IRAK axis as a viable therapeutic target in early clonal hematopoiesis. Collectively, our study expands the landscape of CHIP beyond the canonical mutations. MYD88 mutations, while less common, may significantly impact hematopoietic output and disease risk via sustained immune signaling.

## Acknowledgments

We thank the Comprehensive Mouse and Cancer Core (Jeff Bailey and Victoria Summey), the Viral Vector Core (Thouwa Samake), Flow Cytometry Core, and Genomics Sequencing Core at CCHMC for their assistance. We thank Donald Lynch for helpful discussion and suggestions.

## Data-sharing statement

Cell lines and mouse models used in these studies are publicly available through commercial sources or may be made available from the authors upon written request and material transfer agreement approval. The authors are also glad to share guidance regarding protocols and assays used in these studies upon written request.

## Author Contributions

JY performed experiments, analyzed and interpreted data, and wrote the manuscript. JY, SP, CC, PA, LL, and EU performed experiments and analyzed and interpreted data. KH assisted with the mouse experiments. HCR generated the original Myd88 mutant mice. KC performed bioinformatics analyses. JCB and EH provided input and reagents and interpreted data. JCB, TH, GL, TJD, and AL obtained, designed the targeted CH panel and performed sequencing analysis. PP and KAW analyzed the pathology. DTS conceived and directed the study, analyzed and interpreted data, and wrote and/or edited the manuscript. All authors approved the final version of the manuscript.

## Materials and Methods

### CHIP Patients

Peripheral blood samples were obtained from a cohort of 454 newly diagnosed solid tumor patients from different counties in Appalachia. All patients provided written informed consent for participation in the treatment studies in accordance with the Declaration of Helsinki. DNA extracted from peripheral blood mononuclear cells from patients were analyzed for the presence of CH. Error-corrected next-generation sequencing (NGS) of DNA using the Ion Torrent Personal Genome Machine was used for high definition (HD) sequencing for a detection depth of up to 50,000X (average 44,469X coverage with uniformity of 91.24% and being on target at 99.49%). A custom AmpliSeq HD primers panel IAH150087 was designed to detect CH-2%, which includes whole gene sequencing and hotspots. The region covered in MYD88 is described in **Supplemental Table 7**. DNA was prepped on GeneAmp PCR system 9700 Dual 96-well thermal cycler from Applied Biosystems. PCR products were purified with Agencourt AMPure XP kit (A63881 Beckman Coulter, Indianapolis, Indiana). DNA libraries were prepared with Ion AmpliSeq HD Library kit (A37694) and quantified using real-time PCR with Ion Library TAQMAN Quantitation kit 4468802 on (Applied Biosystems ViiA7 Real-Time PCR System) instrument to allow for an optimal final dilution of the library for template preparation. Bidirectional sequencing with dual barcode support of 454 amplicons in 2 pools at 27.74kb panel size with 99.86% coverage. Template preparation was performed using Ion One Touch2 instrument with Ion 540 Kit OT2 kit (A27753), then enrich and purify Ion One Touch2 ES. Purified ISPs were run on the Ion Torrent S5 instrument using 540 Kit OT2 (A27753) and Ion 540 Chip Kit (A27766). IonAmpliseq HD Dual Barcodes kit (A37695) was used to run multiple samples on the same chip. Data was collected and analyzed on S5 Prime Sequencer Server with Torrent Suite 5.12.2. Final analysis of sequence data was performed using a combination of software: Variant Caller v.5.12.27-1, IGV5.01 (0) and Ion Reporter v.5.18. The hg19 reference sequence was used for manual analysis to assess for deviation from the reference sequence and to evaluate the quality of the sequence and the depth of coverage. Clonal hematopoiesis of MYD88 was defined with a variant allelic frequency (VAF) of 2% or greater.

### Reagents

The inhibitor, NCGC-1481, compound was previously described^63,64^. 4-OHT Hydroxy-tamoxifen and tamoxifen were purchased from Millipore-Sigma. Respective human and murine cytokines, hIL3, hIL6, hTPO, mFLT3, mSCF, and mIL3, were purchased from PeproTech.

### Cell lines

CRISPR/Cas9 targeting human MYD88 (Synthego) was used to generate MYD88 knockouts in parental THP1 and MDSL cell lines^60^. The following sgRNAs supplied by Synthego were used: sgRNA 1: UCCUGGAGCCUCAGCGCGGU; sgRNA 2: GGAGGAUGUGGAGGAGACCG; sgRNA 3: GUUCUUGAACGUGCGGACAC. MYD88 knockout cells were generated by suspending parental THP1 or MDSL cells in buffer R with Cas9-NLS and sgRNA mixture, and electroporated (1700 mV x 20 ms x 1 pulse) using the Neon Transfection system (ThermoFisher). THP1 and MDSL cells were cultured in RPMI 1640 medium supplemented with 10% FBS and 1% penicillin-streptomycin. For MDSL cells, media was supplemented with 10 ng/mL of hIL-3. Primary murine BMNCs (bone marrow monocular cells) were cultured in Iscove’s MDM with 10% FBS, 1% penicillin-streptomycin, and supplemented with 50 ng/mL of hIL6, hTPO, mSCF, mIL3, and mFLT3 cytokines.

### Retroviral vectors

HA-tagged MYD88 plasmid was purchased from Addgene (#12287) and cloned into pCDH-EF1-IRES-copGFP expressing vector with XbaI and BstBI and used as template for MYD88 mutants. MYD88 mutants were generated with Phusion Site-Directed Mutagenesis Kit (F541, ThermoFisher Scientific). The following MYD88 primers were used to generate site-directed mutations using: L265P forward: P-CATCAGAAGCGACCGATCCCCATCAAG; L265P reverse: P-GGCACCTGGAGAGAGGCTGAGTGCAAA; T71I forward: P-ACAAGCGGACCCCATTGGCAGGCTGCT; T71I reverse: P-GTCTCCAGTTGCCGGATCTCCAAGTA; R140Q forward: P-AGCAGTGTCCCACAGACAGCAGAGCTG; R140Q reverse: P-GTCTACAGCGGCCACCTGTAAAGGCTT; C203R forward: P-CTGCCTGGCACCCGTGTCTGGTCTATT; C203R reverse P-GACATCGCGGTCAGACACACACAACTT

### Mice

MYD88^L252P/L252P^ mice (ortholog of the human MYD88 p.L265P mutation), which had been previously generated and described,^37^ were purchased from Jackson Laboratory (Bar Harbor, ME) and bred with Rosa26-CreERT2 mice (Jackson Laboratory) to generate littermates for all subsequent *in vivo* bone marrow transplantations and *in vitro* studies. experiments. All mouse experimental procedures were performed and bred in-house in accordance with and ethically approved by the Institutional Animal Care and Use Committee (IACUC) at Cincinnati Children’s Hospital Medical Center.

### Bone marrow transplantation

For noncompetitive BM transplantations, BMNCs (bone marrow monocular cells) were isolated from femur and tibia of age-matched MYD88^WT/WT^;Rosa26CreERT2 or MYD88^L252P/L252P^;Rosa26CreERT2, grounded using mortar and pestle in sterile 1 XPBS with 1 mM EDTA, and filtered through a 100 μm strainer. Filtered BMNCs were pelleted at 800xg for 10 mins at 4°C, followed by red blood cell lysis in BD Pharm Lyse (BD Biosciences, 555899) and washed with sterile 1X PBS. CD45.2^+^ MYD88^WT/WT^;Rosa26CreERT2 or MYD88^L252P/L252P^;Rosa26CreERT2 BMNCs (1 × 10^6^) were resuspended in 200 μL of PBS and intravenously (i.v.) administered into lethally-irradiated (7.0 Gy and 4.75 Gy after 3 h) recipient mice (CD45.1^+^ B6.^SJLPtprcaPep3b/Boy^ (BoyJ); 6-10 weeks of age) as previously described^15,65,66^. For competitive BM repopulation, BMNCs derived from littermates, age and gender-matched MYD88^WT/WT^;Rosa26CreERT2 or MYD88^L252P/L252P^;Rosa26CreERT2 (CD45.2^+^) femur and tibia, were mixed with equal number of CD45.1^+^ BoyJ BMNCs and then injected i.v. into lethally-irradiated recipient CD45.1 BoyJ. At 4 weeks post-BM transplantation, recipient mice were injected intraperitoneally (I.P.) with 50 μL of tamoxifen (1 mg dissolved in corn oil) daily for 2 weeks to allow tamoxifen-inducible excision of floxed regions and expression of MYD88^L252P/L252P^. For *in vivo* IRAK1/4 inhibitor study, 50 μL of NCGC-1481 at 30 mg/kg or vehicle control (1X PBS) was IP administered daily for 6 weeks.

### Single-cell RNA-sequencing of cKit+ BM cells

BMNCs were isolated from femur and tibia, followed by RBC lysis as describe above, and incubated with cKit-enrichment magnetic antibodies against CD117 (130-091-224, Miltenyi). CD117+ HSCs were positively selected or purified in LS magnetic separation columns (130-042-401, Miltenyi). cKit-enriched cells were then incubated with a panel of cell hashing antibodies (TotalSeq-B, 155831, 155833, 155835, 155837, 155839, 155841, BioLegend) on ice for 30 mins. The scRNA-Seq assay was performed according to the manufacturer’s instructions (Chromium Next GEM Single Cell 3’ Reagent Kits v3.1 (Dual Index) with Feature Barcode technology for Cell Surface Protein, 10x Genomics). Briefly, Total-Seq B antibody-labeled cells were resuspended in the master mix and loaded together with partitioning oil and gel beads into the chip to generate a gel bead-in-emulsion (GEM). The poly-A RNA from the cell lysate contained in every GEM was reverse transcribed into cDNA, adding an Illumina TruSeq R1 primer sequence, Unique Molecular Identifier ^33^ and the 10x Barcode. The DNA conjugated to the Total-Seq B antibodies (Feature Barcodes (FBs)) was barcoded by adding an Illumina Nextera R1, UMI, and the 10x Barcode. The cell barcoded molecules were then cleaned up with Silane DynaBeads and amplified using 13 PCR cycles. Size selection using SPRIselect reagent was performed post amplification to separate full-length cDNA from FBs. Next, full-length, barcoded cDNA was then enzymatically fragmented, sized-selected, adapter-ligated, and amplified for library construction. During the library construction, P5, P7, i7 and i5 sample indexes, and TruSeq Read 2 were added. Separately, FBs were prepared into library constructs by incorporating P5, P7, i7 and i5 sample indexes, and TruSeq Read 2 via PCR. Samples were pooled and run on the NovaSeq X Plus sequencer with a 10B flow cell using the following sequencing parameters: R1: 28 cycles, i7: 10 cycles, i5: 10 cycles, R2: 90 cycles. The 10x Genomics scRNA-Seq libraries from mouse samples were aligned to the mm10 mouse genome and pre-processed using the Cell Ranger-multi) pipeline (v7.2.0, https://www.10xgenomics.com/support/software/cell-ranger) with custom TotalSeq-B hashtag oligos. Normalization, dimensional reduction, clustering, integration and all downstream analyses were performed using Seurat (v5.0.3, https://satijalab.org/seurat/) on R/4.2.3 (https://www.r-project.org). Doublets were detected and removed using DoubleFinder (v2.0.4, https://github.com/chris-mcginnis-ucsf/DoubletFinder) and downstream analyses were done using remaining cells from 900 WT MYD88 900 and 1195 MYD88^L252P/L252P^ cells. For clustering functions, the dimension was set to 1:30 and resolution to 0.8. During sample integration, default parameters were used in Seurat’s FindIntegrationAnchors function, including setting anchor features to 2000. Each cluster was initially annotated using Seurat’s label-transfer function with the HSPC reference atlas ^67^ from Preleukemic Mouse Cell Atlas^68^, then manually re-annotated using top 100 cell marker genes as previously described^69^. The lineage trajectory and pseudo-time were predicted using Monocle3 (v1.3.4, https://cole-trapnell-lab.github.io/monocle3) with resolution = 1^56^.

### Quantitative RT-PCR

Total RNA of cells were isolated using Quick-RNA MiniPrep kit (Zymo Research) and 1 μg of RNA was reverse transcribed into cDNA using High-Capacity RNA-to-cDNA kit (4387406, ThermoFisher Scientific) according to the manufacturer’s procedure. cDNA was diluted 1:5 prior to performing real-time quantitative PCR, using SYBR Green Master Mix (4309155, ThermoFisher Scientific), on an Applied Biosystems StepOne Plus Real-Time PCR System.

### Hematological and histological analysis

Femur or spleen were fixed in formalin and then stained with hematoxylin and eoisin. Complete blood counts of PB was measured by hemacytometer (HEMAVET).

### Flow cytometry analysis

For flow cytometric analysis of lineage positive cells or stem/progenitor HSCs, PB or BM samples were processed in 1xRBC lysis for 15 minutes, followed by incubation in the following antibodies, DAPI (D1306, ThermoFisher Scientific), 7AAD (00-6993-50, eBiosciences), CD11b-PE-Cy7 (25-0112-81, eBiosciences), Gr1-eFluor450 (48-5931-82, eBiosciences), CD3-PE (12-0031-83, eBiosciences), B220-APC (17-0452-82, eBiosciences), CD45.1-Brilliant Violet 510 (110741, BioLegend), CD48-APC (11-0481-85, eBiosciences), CD117-APC-Cy7 (135135, BioLegend), Ly-6A/E(Sca-1)-PE (12-5981-82, eBiosciences), CD135-PE-Cy5 (135312, BioLegend), CD150-PerCpCy5.5 (115922, BioLegend), CD127-Brilliant Violet 605 (35041, BioLegend) and CD45.2-APC-eFluor780 (47-0454-82, eBiosciences) or CD45.2-eFluor450 (48-0454-82, eBiosciences).

### Immunoblot analysis

Mouse BMNCs were collected, RBCs lysed, followed by cell lysis in cold 1XRIPA with protease inhibitors. Protein concentration was quantified by bicinchoninic acid (BCA) assay (32106, Pierce) followed by resuspension in sample loading buffer. Proteins were separated by SDS-PAGE, transferred to nitrocellulose membranes and analyzed by immunoblotting. Immunoblotting was performed with the following primary antibodies: phospho-RelA (3033, Cell Signaling Technology), RelA (sc-71675, Santa Cruz), phospho-IKKα/β (2697, Cell Signaling Technology), IKKβ (2370, Cell Signaling Technology), vinculin (13901, Cell Signaling Technology), phospho-IRAK1 T209 (A1074, Assay Biotech), IRAK1 (sc-5288, Santa Cruz), phospho-IRAK4 T345/S346 (11927S, Cell Signaling Technology), IRAK4 (4363, Cell Signaling Technology), MYD88 (sc-136970, Santa Cruz).

### Lentiviral transduction

As previously described, HEK293T cells were transfected with HA-tagged MYD88 wild-type or mutants and viral packaging (gag-pol and VSV-G) plasmids using TransIT-®LT1 transfection reagent to generate lentiviral supernatants 48 hours post-transfection.^15^ Viral supernatant was harvested and filtered onto isogenic MYD88 KO cells. Transduced cells were expanded and then sorted for GFP+ population.

### Bulk RNA-sequencing

Total RNA of transduced THP-1 MYD88 knockout cells was isolated using Quick-RNA MiniPrep kit (Zymo Research). The initial amplification step for all samples was done with the NuGEN Ovation RNA-Seq System v2. The assay was used to amplify RNA samples to generate cDNA, followed by Qubit concentration quantification. Libraries were then generated using the Illumina protocol (Nextera XT DNA Sample Preparation Kit). The size of the libraries was measured using the Agilent HS DNA chip. The concentration of the pool was optimized to acquire at least 15-20 million reads per sample.

### Gene set enrichment analysis

The analysis of RNA sequencing was performed using iGeak ^70^ and gene set enrichment analysis (GSEA) was performed as previously described^71^.

### Clonogenic progenitor assays

BMNCs were isolated from femur and tibia, followed by RBC lysis and incubation with cKit-enrichment magnetic antibodies against CD117 and positively selected or purified in LS magnetic separation columns. CD117^+^ HSCs were resuspended in Iscoves MDM (10-016-CV, Corning), and 1000 cKit^+^ BM were treated with 1 μM 4-OHT and plated in MethoCult GF (M3434 or M3534; StemCell Technologies). Colonies were enumerated using StemVision (StemCell Technologies) on days 12-14. To assess serial replating capacity, colonies were pooled, resuspended in Iscoves MDM, washed twice, and then replated serially at 7.5×10^3^ cells per replicate in the same medium.

### Profiling of plasma cytokines and chemokines by multiplex ELISA

Peripheral blood (PB) was collected in EDTA tubes from the indicated non-competitive BMT recipient mice at 12 weeks post-BMT. PB was centrifuged at 1000xg for 10 mins at 4°C for plasma in the supernatant. The supernatant was transferred to cold Eppendorf tubes and immediately frozen at −70°C. Samples were thawed on ice and mixed thoroughly before being diluted 1:1 in assay buffer provided by the mouse cytokine/chemokine magnetic bead panel kit (MCYTMAG-70K-PX32, Millipore). Luminex xMAP platform was used to quantify the 32-plex mouse cytokine panel.

### Data Availability

The RNA sequencing data generated in this study has been deposited at NCBI’s GEO repository with accession number GSE277714 (token: qzylicsqhzujpid).

### Statistical analysis

Differences among multiple groups were assessed by one-way analysis of variance (ANOVA) followed by Tukey’s multiple comparison posttest for all possible combinations. Comparison of two groups was performed using the Mann-Whitney test or the Student’s t test (unpaired, two-tailed) when sample size allowed. Unless otherwise specified, results are depicted as the mean ± standard deviation or standard error of the mean (SEM). A normal distribution of data was assessed for data sets >30. For correlation analysis, Pearson correlation coefficient (r) was calculated. For Kaplan-Meier analysis, Mantel-Cox test was used. All graphs and analyses were generated using GraphPad Prism 10 software or using the package ggplot2 from R.

---

**Supplemental Figure 1.**
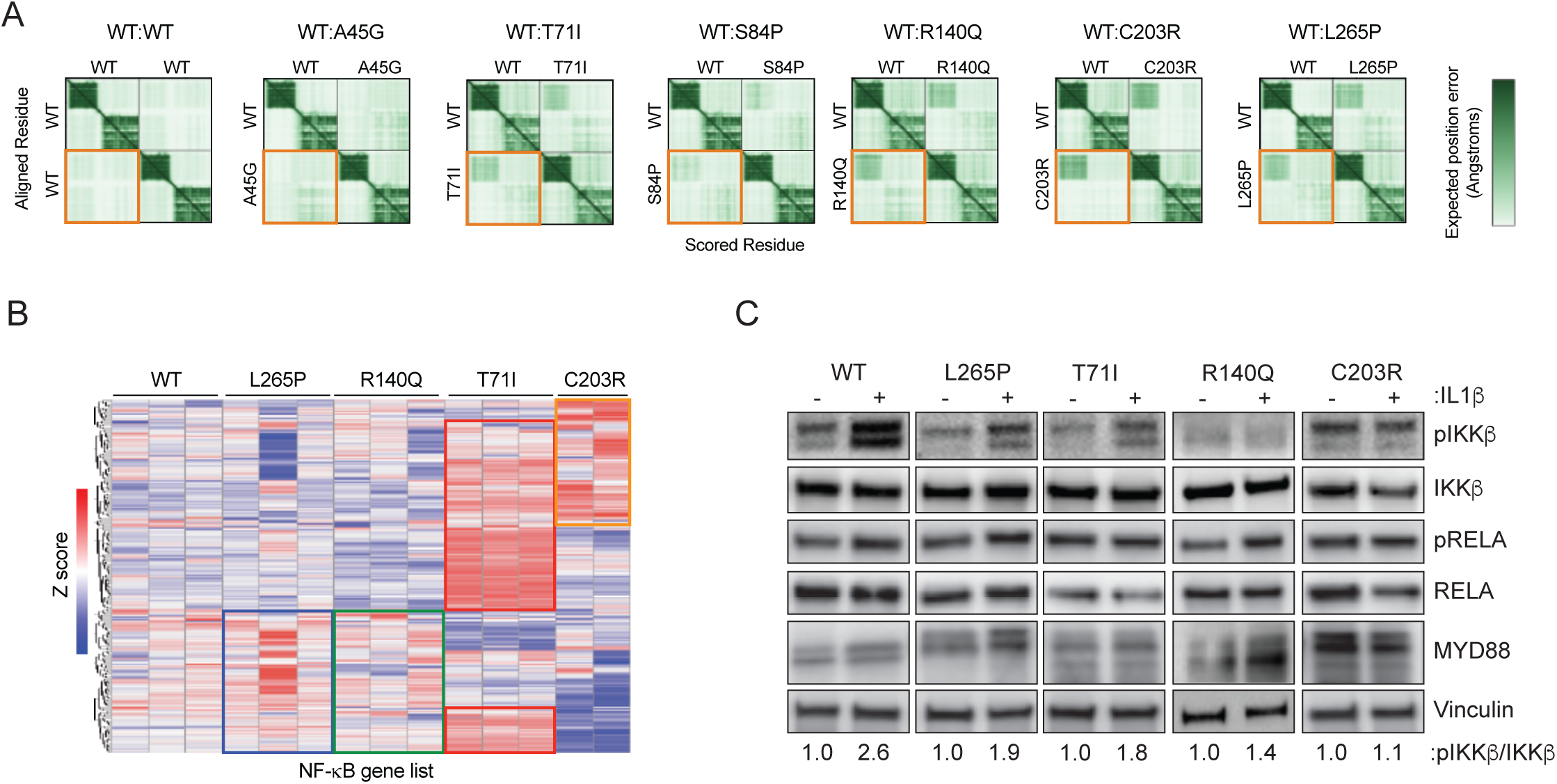
Characterization of MYD88 CHIP mutant. **(A)** Representative PAE analysis of human MYD88 dimers. The lower left quadrant (orange box) shows the expected possition error for the respective dimers: WT and WT, WT and A45G, WT and T71I, WT and S84P, WT and R140Q, WT and C203R, and WT and L265P. **(B)** Heatmap of differentially expressed NF-κB target genes (Z score) from THP1 MYD88 KO cells transduced with lentiviral vectors encoding WT MYD88 or the CHIP mutations (L265P, T71I, R140Q, and C203R). **(C)** Immunoblot analysis of THP1 MYD88 KO cells transduced with lentiviral vectors encoding WT MYD88 or the CHIP mutations (L265P, T71I, R140Q, and C203R) follwoing treatment with IL-1β. Ratio of pIKKβ relative to total IKKβ is shown below.

**Supplemental Figure 2.**
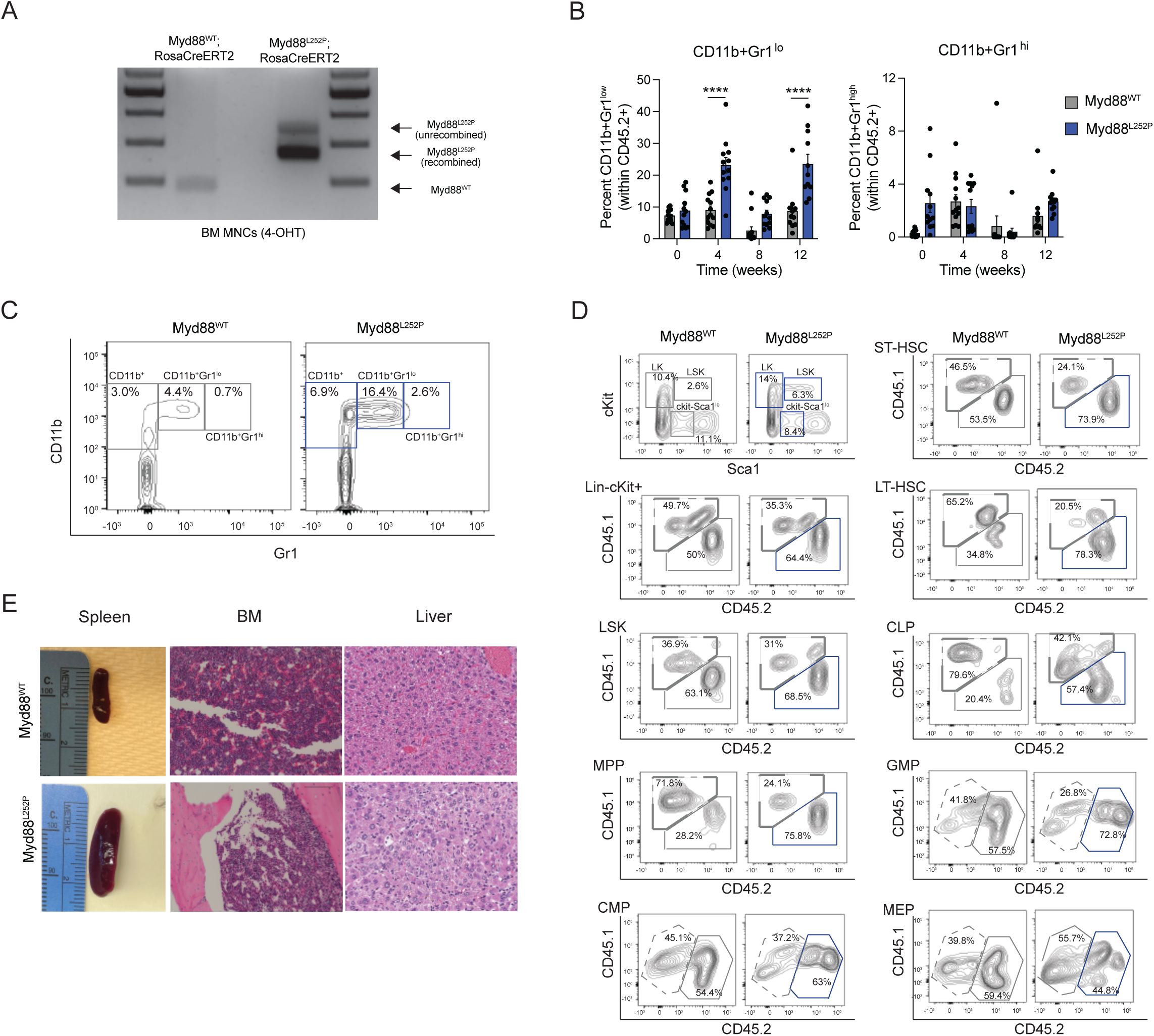
Characterization of Myd88^L252P^ mice. **(A)** Genotyping analysis of Myd88^WT^;Rosa-CreERT2 and MyD88^L252P^;RosaCreERT2 homozygous mice following 4-OHT induced recombiation. (**B**) Proportions of donor-derived CD45.2 CD11b+Gr1lo and CD11b+Grhi cells in the PB of Myd88^WT^ and Myd88^L252P^ on left panel. Error bars represent SEM (n = 11-13 per group). **(C)** Representative flow cytometry gating of donor-derived CD45.2 CD11b+Gr1lo and CD11b+Grhi cells in the PB of Myd88^WT^ or Myd88^L252P^. Error bars represent SEM. Significance was determine with a Student’s t-test for two groups or ANOVA for multiple groups (*, P < 0.05; **, P < 0.01; ***, P < 0.001). **(D)** Representative flow cytometry gating of donor-derived CD45.1 and CD45.2 LK (Lin-cKit+Sca1-), LSK (Lin-ckit+Sca1+), LT-HSC (LSK CD150+CD48-), ST-HSC (LSK CD150-CD48-), MPP (LSK CD150-CD48+), CMP (LK CD34+16/32-), MEP (LK CD34-CD16/32-), GMP (LK CD34+CD16/32+), CLP (Lin-ckit-Sca1loCD127+CD135+) from the BM of mice competitively engafted with Myd88^WT^ or Myd88^L252P^ BM cells. **(E)** Representative images of spleens and BM sections (Hematoxylin and Eosin) from recipient mice transplanted with Myd88^WT^ and Myd88^L252P^ BM cells.

**Supplemental Figure 3.**
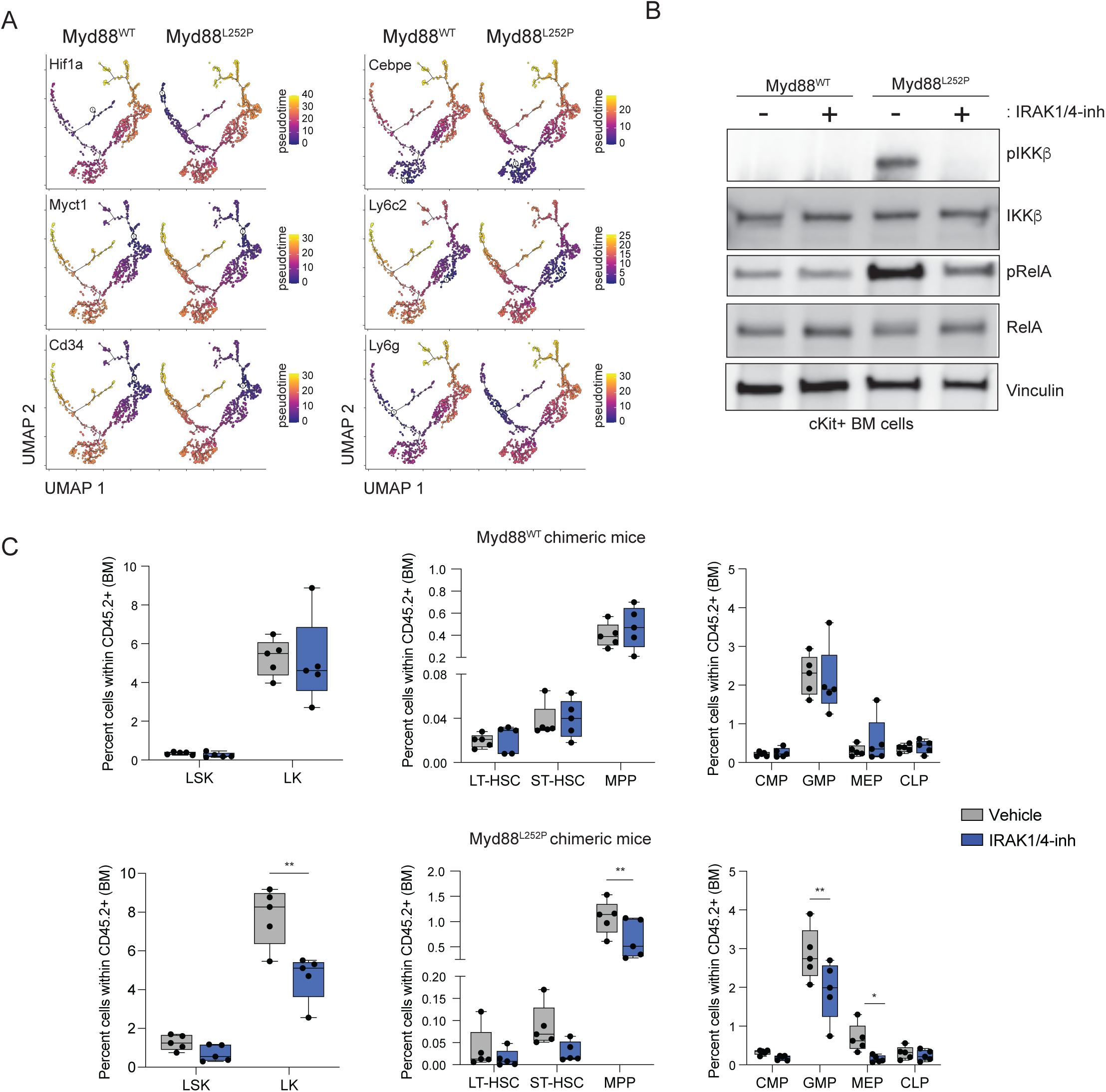
Pseudotime analysis and IRAK1/4-inhibitor treatment of Myd88^L252P^ BM cells. **(A)** BM cells were harvested, and cKit+ cells were enriched by magnetic selection. Cells were analyzed by 10X single-cell RNA (scRNA) sequencing. UMAP of 10,000 cells representing 13 clusters from recipient mice transplanted with Myd88^WT^ or Myd88^L252P^ BM collected at 12 weeks. Pseudotime trajectory of scRNA-seq profiles of the major clusters as defined by the indicated genes. **(C)** Immunoblotting of Myd88^WT^ and Myd88^L252P^ cKit+ BM cells incubated with 4-OHT to induce recombination for 72 hours followed by treatment with vehicle (DMSO) or 1 μM IRAK1/4-inhibitor (IRAK1/4-inh; NCGC-1481) for 24 hours. **(C)** Proportion of donor-derived Myd88^WT^ (n = 5) and **(C)** Myd88^L252P^ (n = 5) CD45.2 populations from the BM of cBMTs following treatment with vehicle or IRAK1/4-inh: LK (Lin-cKit+Sca1-), LSK (Lin-ckit+Sca1+), LT-HSC (LSK CD150+CD48-), ST-HSC (LSK CD150-CD48-), MPP (LSK CD150-CD48+), CMP (LK CD34+16/32-), MEP (LK CD34-CD16/32-), GMP (LK CD34+CD16/32+), CLP (Lin-ckit-Sca1loCD127+CD135+). Data is representative from 2 independent biological replicates. Error bars represent SEM. Significance was determined with a Student’s t-test for two groups or ANOVA for multiple groups (*, P<.05; **, P < 0.01).

